# MetaSanity: An integrated, customizable microbial genome evaluation and annotation pipeline

**DOI:** 10.1101/789024

**Authors:** Christopher J Neely, Elaina D Graham, Benjamin J Tully

## Abstract

**Summary:** As the importance of microbiome research continues to become more prevalent and essential to understanding a wide variety of ecosystems (e.g., marine, built, host-associated, etc.), there is a need for researchers to be able to perform highly reproducible and quality analysis of microbial genomes. MetaSanity incorporates analyses from eleven existing and widely used genome evaluation and annotation suites into a single, distributable workflow, thereby decreasing the workload of microbiologists by allowing for a flexible, expansive data analysis pipeline. MetaSanity has been designed to provide separate, reproducible workflows, that (1) can determine the overall quality of a microbial genome, while providing a putative phylogenetic assignment, and (2) can assign structural and functional gene annotations with varying degrees of specificity to suit the needs of the researcher. The software suite combines the results from several tools to provide broad insights into overall metabolic function and putative extracellular localization of peptidases and carbohydrate-active enzymes. Importantly, this software provides built-in optimization for “big data” analysis by storing all relevant outputs in an SQL database, allowing users to query all the results for the elements that will most impact their research.

**Availability and implementation:** MetaSanity is provided under the GNU General Public License v.3.0 and is available for download at https://github.com/cjneely10/MetaSanity. This application is distributed as a Docker image. MetaSanity is implemented in Python3/Cython and C++.

**Supplementary information:** Supplementary data are available below.

## 1 Introduction

The analysis of microbial genomes has become an increasingly common task for many fields of biology and geochemistry. Researchers can routinely generate hundreds/thousands of environmentally derived microbial genomes using methodologies such as metagenomics (Tully et al., 2018), high-throughput culturing (Thrash et al., 2015), and single cell sorting (Stepanauskas et al., 2017). However, analyzing the data can be problematic, as data analysis is computationally intensive and requires a knowledge of software that is constantly changing and may be difficult to install or execute. For the average researcher, the task of evaluating and annotating a set of microbial genomes may be time intensive and computationally rigorous. Here, we present MetaSanity, a comprehensive and customizable solution for generating evaluation and annotation pipelines for bacterial and archaeal isolate genomes, metagenome-assembled genomes (MAGs), and single-amplified genomes (SAGs). MetaSanity provides genome quality evaluation, phylogenetic assignment, as well as structural and functional annotation through a variety of integrated programs based on the procedure described in ref. Tully (2019). MetaSanity provides a workflow that combines all outputs into a single queryable database that operates easily from the command line. Installation can be performed at the user level, limiting the need for intervention by system administrators, and, except for certain memory intensive programs, can be run locally on high-end personal computers.

## 2 Description of Methods

MetaSanity consists of two smaller workflows (Figure 1): (1) PhyloSanity, to evaluate the completion, contamination, redundancy, and phylogeny of each genome in a dataset, and (2) FuncSanity, to provide structural and functional annotations of each genome. Each component consists of several optional applications that can be customized to specific research needs. While each component contained within the two pipelines runs independently and generates component specific outputs, MetaSanity combines all outputs into a single queryable SQL database that allows fast and easy retrieval of data – in this case, gene annotations and other related genomic data. MetaSanity focuses on allowing users the ability to fine-tune and customize their data analysis pipelines with minimal effort and maximized computational and storage efficiency (Supplemental Table 1). MetaSanity is distributed as a Docker image (Merkel et al., 2014) and is implemented using a combination of Python3 (Python Software Foundation 2014) /Cython (Bradshaw et al., 2011) and C++ (ISO/IEC 2014).

**Figure 1.**
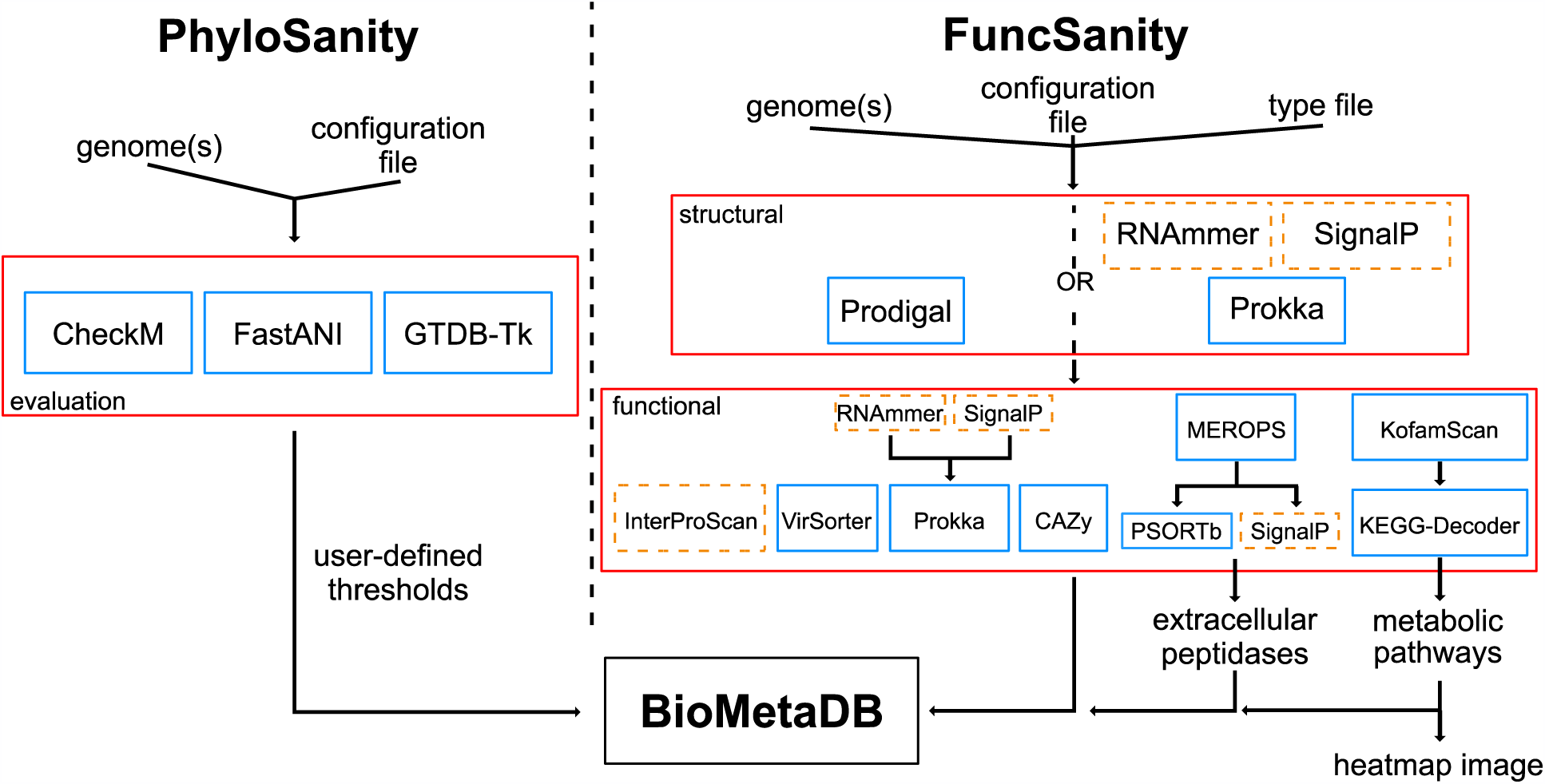
MetaSanity pipeline schema. Programs and databases that are part of the MetaSanity installation are in blue boxes. Programs in the dotted orange boxes must be installed separately by the user due to licensing agreements.

### 2.1 PhyloSanity

PhyloSanity is designed to provide metrics of genome quality and to filter genomes for downstream analysis based on user defined quality metrics. The workflow integrates CheckM v1.0.18 (Parks et al., 2015), GTDB-Tk v.0.3.2 (Parks et al., 2018), and FastANI (Jain et al., 2019) as part of its evaluation pipeline. CheckM estimates the completion and contamination of each genome (Parks et al., 2015). Next, FastANI compares each genome in a pairwise fashion against all other genomes to determine the average nucleotide identify (ANI) for each genome pair (Jain et al., 2019). For any set of genomes that shares an ANI above a user-defined value, a non-redundant genome representative will be selected from the set that is the most complete and least contaminated. This allows users the option to exclude redundant genomes from further analysis. Differentiating genomes as non-redundant versus redundant can be useful for researchers working with MAGs or SAGs that are generated from replicate samples and may not have biological meaning when working with isolates or strain level differences. All genomes can undergo phylogenetic assignment based on relative evolutionary distance (Parks et al., 2018) through GTDB-Tk, which will replace the CheckM-returned taxonomic assignment.

### 2.2 FuncSanity

FuncSanity provides structural and functional annotation of microbial genomes. The workflow incorporates annotation suites from eight existing and widely used programs. The use of multiple annotation programs has the advantage of capturing functional predictions that may not have been detected due to database or search limitations. Specialized annotation programs, such as VirSorter (Roux et al., 2015), use custom tools and/or databases to return relevant annotations that are not captured by other programs in MetaSanity. Open reading frames (ORFs) are predicted using Prodigal v2.6.3 (Hyatt et al., 2010); however, users may opt to use the putative coding DNA sequences (CDS) generated by Prokka v1.13.3 (Seeman, 2014). From here, putative ORFs are processed by a set of annotation tools that can be selected by the user with user-defined filtering and cutoff values.

#### Kyoto Encyclopedia of Genes and Genomes (KEGG) Annotation

Putative ORFs can be searched against the KofamKOALA database using KofamScan v.1.1.0 (Aramaki et al., 2019). Default parameters are used and the ‘mapper’ tab-delimited output option is generated, linking ORF IDs to KEGG Ontology (KO) IDs. Users can query any KO ID to generate specific functional search results in BioMetaDB.

#### KEGG-Decoder

KEGG annotations can be used to estimate the completeness of various biogeochemically-relevant metabolic pathways in a genome using KEGG-Decoder v.1.0.1 (Graham et al., 2018; https://github.com/bjtully/BioData/tree/master/KEGGDecoder). Users can search genomes based on completeness of a pathway or function of interest. An additional heatmap summary visualization is generated.

#### VirSorter

VirSorter v1.0.5 (Roux et al., 2015) can be implemented to identify phage and prophage signatures in each genome using default parameters. Users can search for matches to each of the phage and prophage categories returned by VirSorter and generate lists of contigs and/or genomes with the assignments (Supplementary Table 1).

#### InterProScan

InterProScan 5.36-75.0 (Jones et al., 2019) is an optional installation and can be used for domain prediction on putative ORFs. Users have the option of downloading all of the InterProScan databases, including TIGRfam (Haft et al., 2003), Pfam (Finn et al., 2016), CDD (Marchler-Bauer et al., 2017), and PANTHER (Mi et al., 2019). Each InterProScan database result is indexed separately in BioMetaDB and can be used to return matching genomes using database specific IDs (e.g., PF01036 would return putative rhodopsin ORFs from a Pfam result).

#### Prokka Annotation

If not chosen as the option for structural annotation, genomes can be annotated using Prokka and its associated databases with the parameters --addgenes (adds the “gene” feature to each CDS in the GenBank output format), --addmrna (adds the “mRNA” feature to each CDS in the GenBank output format), --usegenus (use the genus-specific databases), --metagenome (improve gene predictions for fragmented genomes), and --rnammer (sets RNAmmer as the preferred rRNA prediction tool). rRNA identification with RNAmmer v.1.2 (Lagesen et al., 2007) and signal peptide detection with SignalP v.4.1 (Nielson, 2017) are optional installations.

#### Carbohydrate-active enzyme (CAZy) Annotation

Putative ORFs can be assigned a putative CAZy functionality (Cantarel et al., 2009) based on the dbCANv2 database (Zhang et al., 2018). ORFs are searched against dbCANv2 using HMMER v3.1b2 (Eddy, 2011) with the minimum score threshold set to 75 (-T parameter). PSORTb v.3.0 (Yu et al., 2010) and SignalP can be optionally performed on CAZy matches to determine if a putative enzyme is predicted to be extracellular. An extracellular assignment is made if PSORTb predicts “extracellular” or “outer membrane” localization or if PSORTb returns “unknown” localization and SignalP predicts the presence of a signal peptide. Users can search for genes and genomes based on overall CAZy annotations or by searching for specific designations (e.g., GT41 for glycosyl transferase family 41).

#### Peptidase Annotation

Putative ORFs can be assigned to a peptidase family using a set of HMMs that represent the MEROPS database (Rawlings et al., 2013). The putative extracellular nature of a MEROPS match can be determined as above. Users can search for genes and genomes based on overall MEROPS annotations or by searching specific peptidase families.

InterProScan, SignalP, and RNAmmer are not automatically distributed with MetaSanity and require users to download their binaries separately and agree to their individual license requirements.

### 2.3 BioMetaDB

BioMetaDB is a specialized relational database management system project that integrates modularized storage and retrieval of FASTA records with the metadata describing them. This application uses tab-delimited data files to generate table relation schemas via Python3. Based on SQLAlchemy v.1.3.7 (Bayer, 2012), BioMetaDB allows researchers to efficiently manage data from the command line by providing operations that include: (1) the ability to store information from any valid tab-delimited data file and to quickly retrieve FASTA records or annotations related to these datasets by using SQL-optimized command-line queries; and, (2) the ability to run all CRUD operations (create, read, update, delete) from the command line and from python scripts. Output from both workflows is stored into a BioMetaDB project, providing users a simple interface to comprehensively examine their data (Supplemental Table 2). Users can query application results used across the entire genome set for specific information that is relevant to their research, allowing the potential to screen genomes based on returned taxonomy, quality, annotation, putative metabolic function, or any combination thereof.

## 3 Results

MetaSanity was tested on two separate systems – a personal computer with an Intel core i5-4570 CPU @ 3.20 GHz processor with 4 cores and 32 GB of RAM operating the Deepin 15.11 Linux distribution, and an academic server with an Intel Xeon E7-4850 v2 @ 2.30 GHz processor with 96 cores and 1 TB of RAM operating the Ubuntu 18.04.3 LTS Linux distribution. Reduced options were calculated on the personal computer using all four available threads and preset parameter flags that skip memory intensive processes. Complete options were calculated on the academic server using 10 threads and no parameters to reduce memory usage. Runtime results are available in Supplemental Table 3. The current architecture relies on sequential completion of time intensive processes, several of which are optional for users. Ongoing modifications that take advantage of parallelizing these processes should decrease the overall computation time.

---

**Supplementary Table 1.**
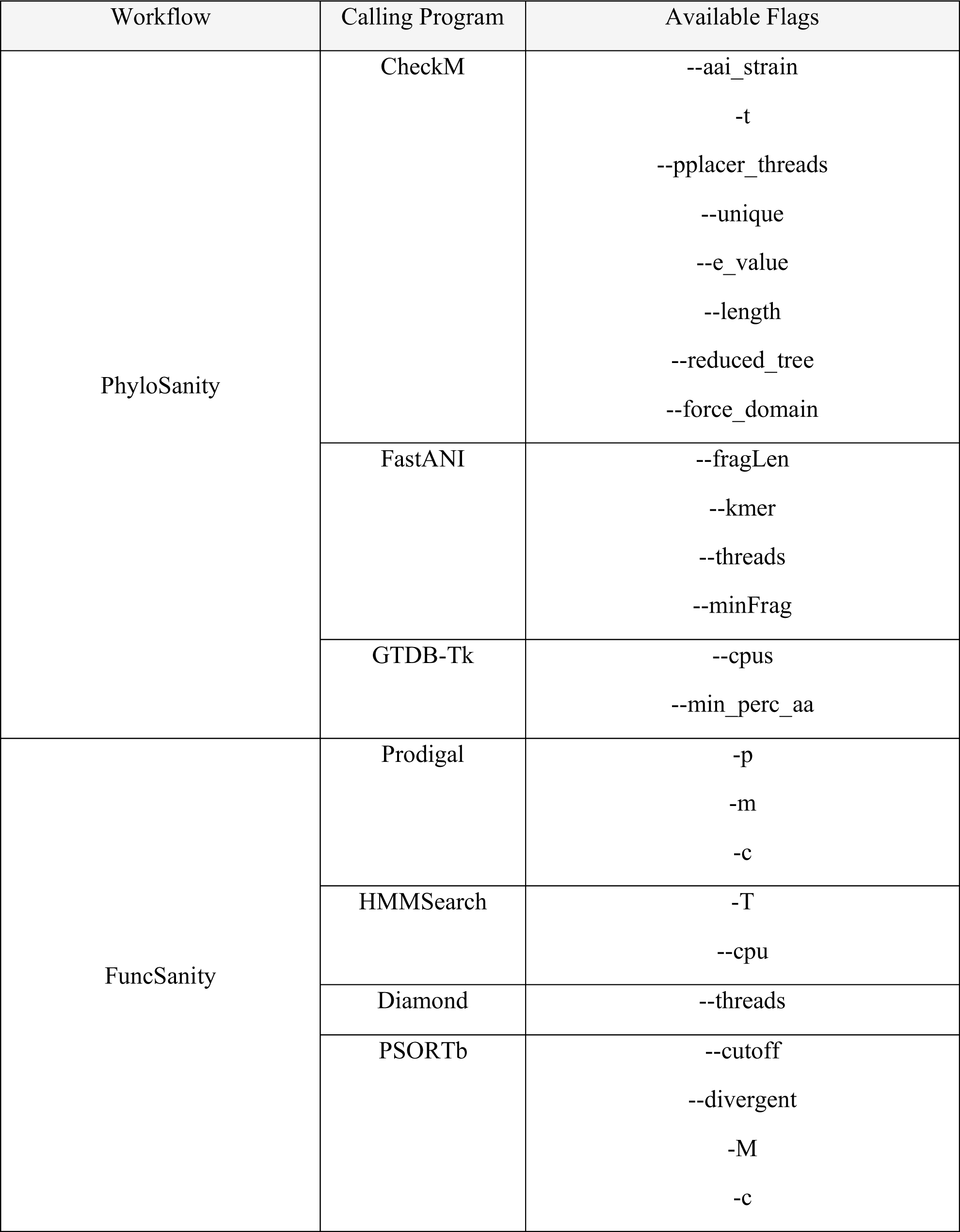
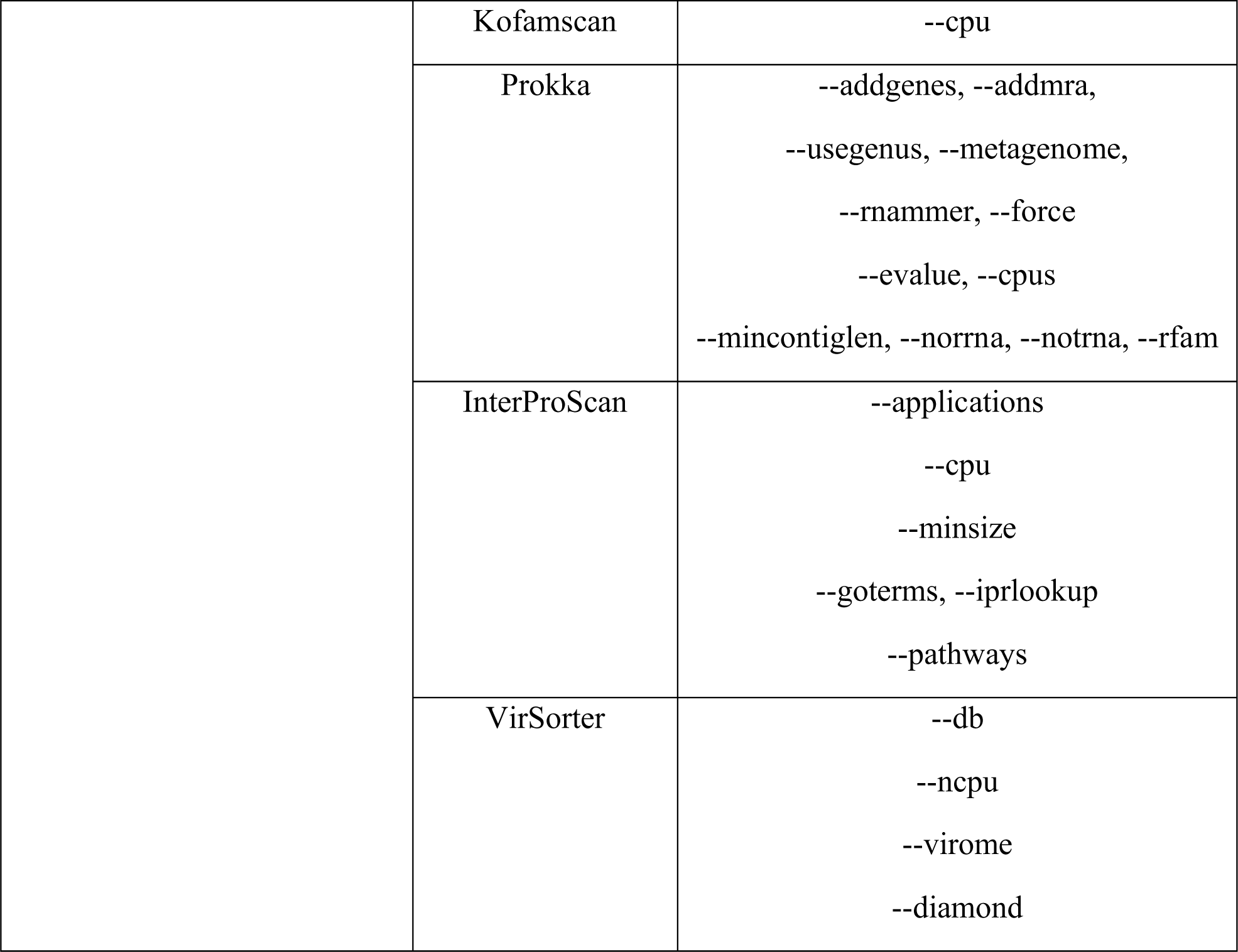
Parameters flags available to each program within the MetaSanity workflows. Modification to the configuration files will allow users may include any set of these flags for specific analyses.

**Supplementary Table 2.**
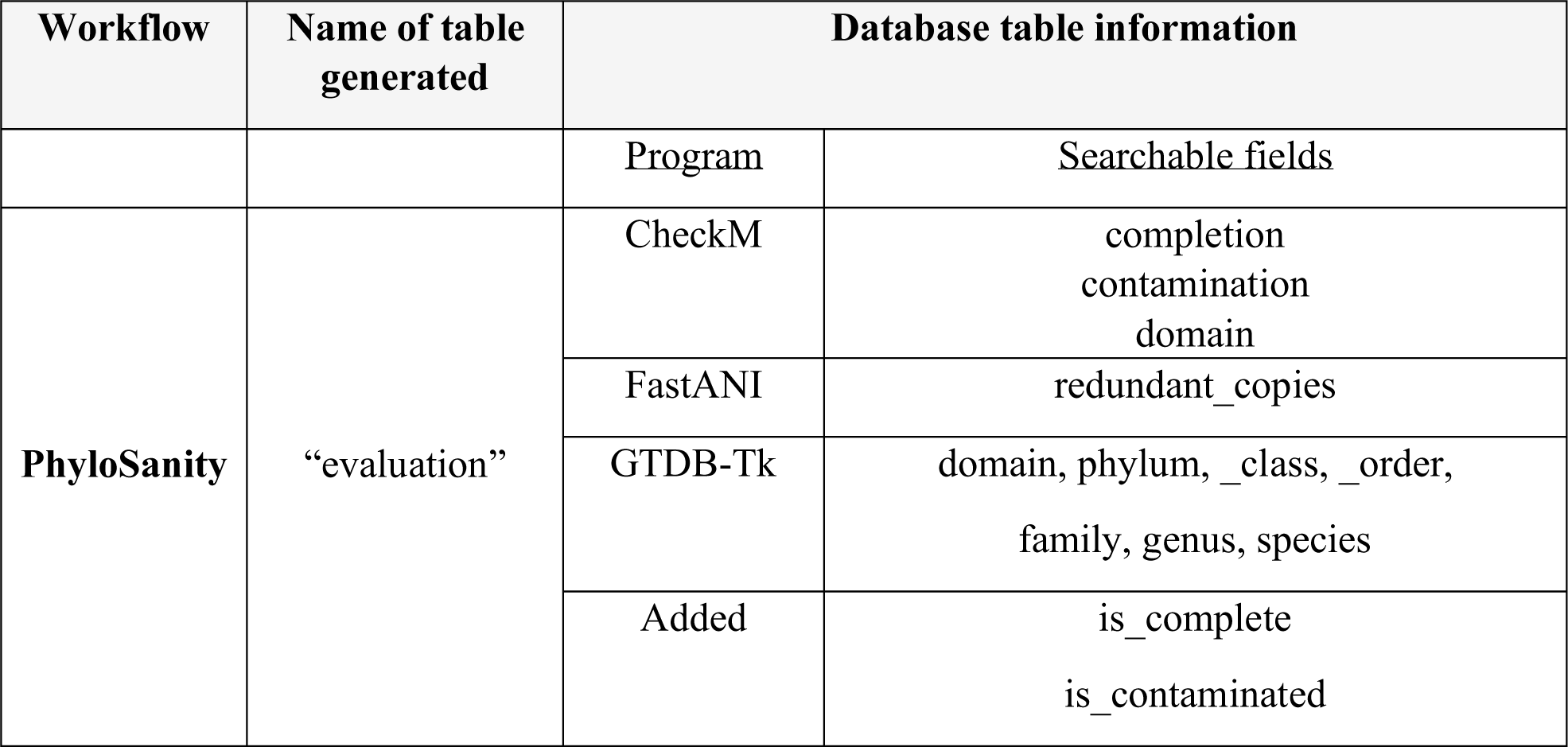
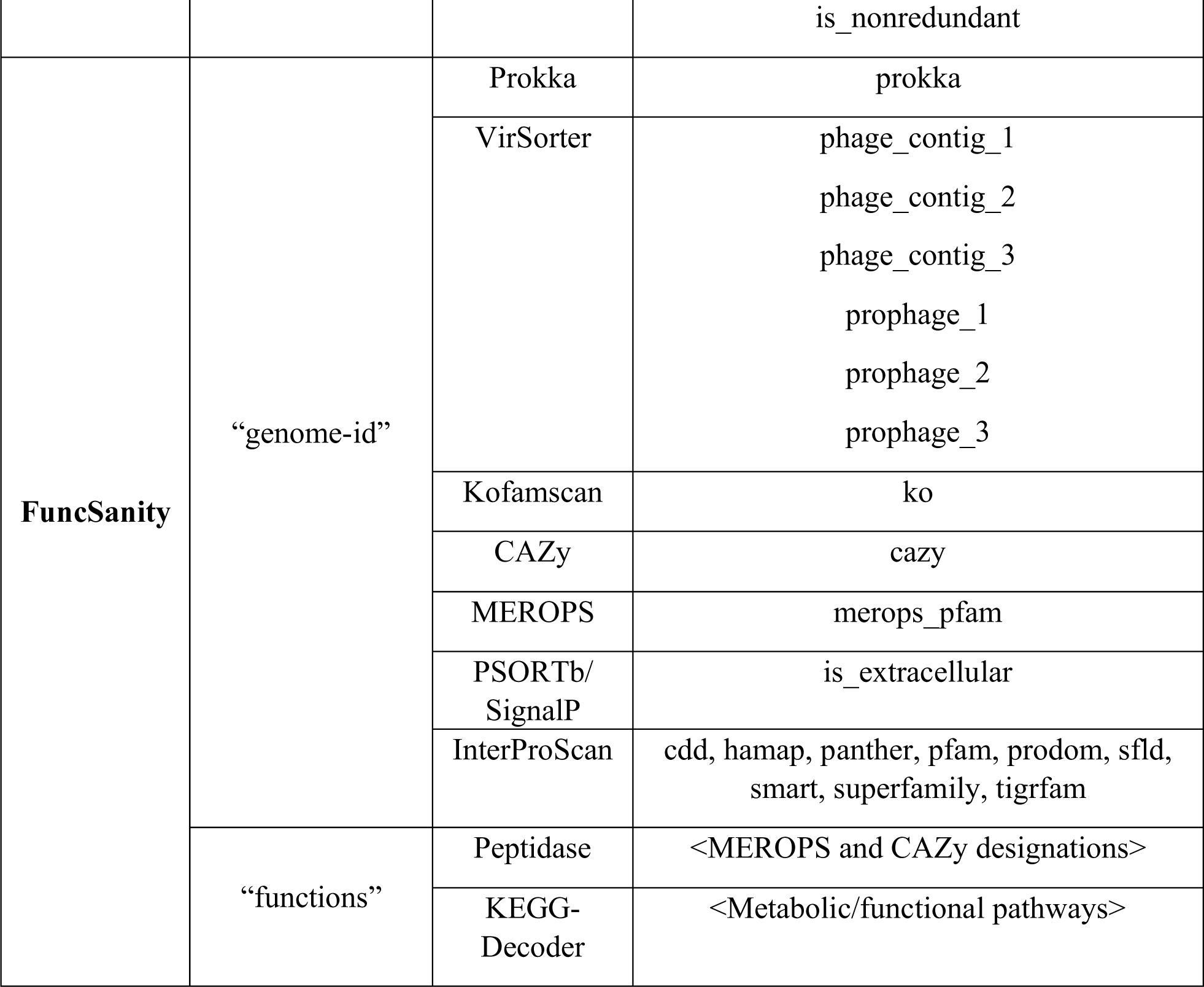
Underlying database structure and queryable objects from the complete MetaSanity workflow.

**Supplemental Table 3.** Runtimes of each core component of the MetaSanity pipeline. InterProScan search ran using the databasesTIGRFAM, SFLD, SMART, SUPERFAMILY, Pfam, ProDom, Hamap, CDD, and PANTHER, with parameter flags --goterms, --iprlookup, and --pathways.

|  | **PhyloSanity** |  | **FuncSanity** |  |
| --- | --- | --- | --- | --- |
|  | No GTDB-Tk;<br>CheckM<br>reduced_tree | Complete<br>evaluation | Recommended<br>& optional; no<br>InterProScan | Complete<br>annotation |
| **1 genome** | 0.05 hr | 0.63 hr | 0.52 hr | 1.62 hr |
| **10 genomes** | 0.15 hr | 1.02 hr | 6.17 hr | 20.3 hr |

**Supplemental Figure 1.**
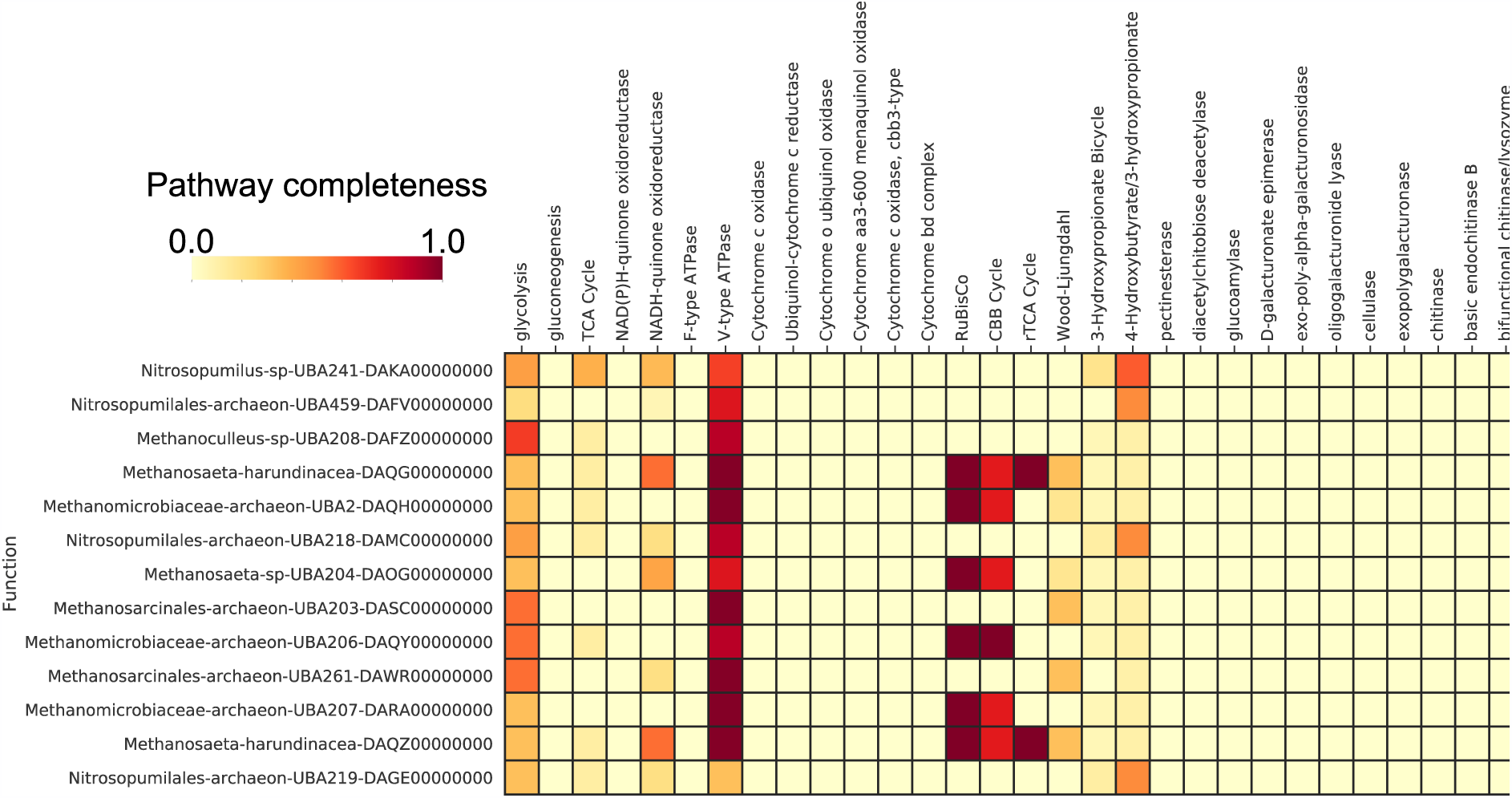
Example KEGG-Decoder heatmap output. The completeness of various biogeochemically-relevant pathways for a collection of marine metagenome-assembled genomes, scaled from 0.0-1.0.

## Acknowledgements

We would like to thank Taylor Reiter, Roth Conrad, Jay Osvatic, and Luiz Irber for providing code contributions to KEGG-Decoder as part of the Moore Foundation funded ‘Speeding Up Science’ hackathon. This is C-DEBI Contribution XXX.

